# Assessment of levamisole HCl and thymosin α1 in two mouse models of amyotrophic lateral sclerosis

**DOI:** 10.1101/2023.01.23.525258

**Authors:** David R. Borchelt, Siobhan Ellison

**Affiliations:** Department of Neuroscience, University of Florida, College of Medicine, Gainesville, FL 32610; Neurodegenerative Disease Research Inc., Fairfield, FL 32634

## Abstract

Amyotrohpic lateral sclerosis (ALS) is a progressive neurodegenerative disease that causes generalized muscle weakness and atrophy. Neuropathologically, ALS is defined by severe loss of upper and lower motor neurons with a robust neuroinflammatory response. In the present study, we have examined the potential utility of two drugs that have indications as immune modulators, levamisole HCl and thymosin α1. These drugs were tested in two models models that reproduce aspects of ALS. We conducted a 14 week dosing study of these two drugs in the SOD1^G93A^ and Prp-TDP43^A315T^ models of ALS. The drugs were given once daily for two weeks and then every other day for 6 weeks for a total of 8 weeks of treatment. Outcome measurements included efficacy assessment on the neuromuscular phenotypes, and pathological analyses of ubiquitin load and neuro-inflammatory markers in spinal motor neurons. Neither of these drug treatments produced significant extensions in survival; however, there were changes in ubiquitin load in SOD1^G93A^ mice that suggest the drugs could be beneficial as additions to other therapies.

## Introduction

Amyotrophic lateral sclerosis (ALS) is a complex disease genetically that is unified by a common theme of upper and lower motor neuron degeneration [1]. Most cases of ALS, both sporadic and familial, display a convergent TDP-43 pathology [2]. Although there has been incremental progress in identifying new therapies for ALS, there is still a great need for therapies that provide profound improvements in life expectancy and quality. Importantly, successful treatment of ALS may require poly-therapies that impact multiple pathways or pathological mechanisms. In the search for new drugs, critical issues for testing the efficacy of treatments are knowledge of the mechanisms or pathways one wants to impact, as well as a model to test the effectiveness of treatment on the target of engagement.

Several ALS rodent models have been used in testing therapies [3]. The SOD1 G93A transgenic mouse was the first model to be generated and made widely available [4]; and thus, is the model that has been used in the majority of pre-clinical drug studies for ALS. The G93A model expresses the mutant human SOD1 gene (harboring a single amino acid substitution of glycine to alanine at codon 93) driven by the human SOD1 promoter [4]. This model shows a mortality/aging phenotype with 50% survival at 128.9+/− 9.1 days and die within 7-14 days after the first signs of weakness [5]. Similar to the G93A model, mice that are transgenic for human TDP43-A315T show abbreviated lifespans [6]. The TDP43 model shows ubiquitin inclusions in some motor neurons with modest motor neuron loss [6]. The G93A SOD1 model is a reliable mimic of motor neuron disease but the TDP43-A315T mice are confounded by gastrointestinal dysfunction that is largely driving abbreviated lifespan [7,8].

In the current study, we investigated the potential utility to two drugs that are already in use in other indications. Thymalfasin (Zadaxin^®^) is a synthetic analog of thymosin α1, which is a 28 amino acid hormone. Thymosin α1 is used in humans as a treatment for hepatitis B, hepatitis C, and some types of cancer. It has been characterized as an immune stimulant that heightens innate immune function [9]. Levamisole HCl (Neuroquel^®^) is also an immune modulating drug that currently is not in use in the United States in humans, but is widely used in veterinary applications. It is used outside the U.S. in humans and has been used as an adulterating agent in illicit street drugs [10]. Levamisole has been used in an ALS clinical trial in which the drug was administered once a week for 6 months; no benefit was observed [11]. Here, we have conducted a 14 week dosing study of these two drugs in the SOD1^G93A^ and Prp-TDP43^A315T^ models of ALS. Levamisole HCl was given once daily for two weeks and then every other day for 6 weeks for a total of 8 weeks of treatment. Thymosin α1 was given subcutaneously twice a week for a total of 8 weeks. Outcome measurements included efficacy assessment on the neuromuscular phenotypes, and pathological analyses of ubiquitin load and neuro-inflammatory markers in spinal motor neurons. Neither of these drug treatments produced significant extensions in survival; however, there were changes in ubiquitin load that suggest the drugs could be beneficial as additions to other therapies.

## Methods

This study was conducted entirely by the Jackson Laboratories (Bar Harbor, ME) under a contract funded by Neurodegenerative Disease Research, Inc. The figures provided are derived from materials provided in final reports by the Jackson Laboratories. Modifications to the figures were limited to changes in the location of markings to indicate statistical differences.

### Experimental Animals

Twelve (12) wild-type and seventy two (72) hemizygous B6SJL-Tg(SOD1*G93A)1Gur/J male mice (JAX stock# 2726) were transferred to the *in vivo* research laboratory in Bar Harbor, ME. Table 1 lists abnormalities that have been described in the G93A model. Twelve (12) wild-type and seventy two (72) hemizygous B6.Cg-Tg(Prnp-TARDBP*A315T)95Balo/J male mice (JAX stock# 010700) were generated by breeding at Jackson Laboratories *in vivo* facility, over 24 weeks. Table 2 lists abnormalities described in TDP43-A315T mice. All studies were conducted by staff at the Jackson Laboratories following protocols approved by the Jackson Laboratories Institutional Animal Use and Care Committee.

**Table 1.** Tissue and phenotype abnormalities exhibited by the SOD1G93A mouse in this study.

|  |  |
| --- | --- |
| Adipose tissue | Reduced adipose tissue accumulation |
| Cardiovascular tissue | Abnormal capillary and vascular endothelial cell morphology |
| Growth and body mass | Lower body mass, decreased weight by 77 days |
| Homeostasis/metabolism | Increased energy expenditure at rest; decreased circulating insulin, decreased plasma leptin |
| Nervous system | Abnormal CNS synaptic transmission, abnormal motor neuron morphology |
| Abnormal gait, locomotor behavior | Abnormal gait at 90 days |
| Cell physiology | Decreased glucose uptake; increased lipid peroxidation, increased sensitivity to oxidative stress |
| Hematopoietic system | Microgliosis |
| Muscle System | Decreased motor units, muscle weakness, muscle atrophy |

**Table 2.** Tissue and phenotype abnormalities exhibited by the TDP43.

|  |  |
| --- | --- |
| Growth and body mass | Weight loss becomes significant at 85 days |
| Mortality and aging | Death occurs at a median of 108 days |
| Homeostasis/metabolism | These phenotypes are associated with gut pathology |
| Nervous system phenotype | Decreased axons (femoral nerve); reactive astrocytosis from 80 days, decreased AchE staining in ganglion of myenteric plexus of the colon; some motor neurons have ubiquitin-positive cytoplasmic inclusions |

The mice were ear notched for identification and housed in individually and positively ventilated polysulfonate cages with HEPA filtered air at a density of 3-4 mice per cage. The animal room was lighted entirely with artificial fluorescent lighting, with a controlled 12 h light/dark cycle (6 am to 6 pm light). Filtered tap water, acidified to a pH of 2.5 to 3.0, and normal rodent chow was provided ad libitum. At 8±1 weeks of age, one hundred and sixty eight (168) mice were grouped by body weight into ten groups. Enrollment was staggered in three cohorts of an equal number of mice in each group, with a 4-6 week intervals between successive cohorts. In several instances, premature death due to self-mutilation necessitated removing a mouse from the survival cohort into a treatment cohort. Tables 1 and 2 list abnormalities that have been described in these two mouse models.

### Treatment

Mice were dosed by gastric gavage (PO) or subcutaneous (SQ) injection, for up to 8 weeks, as described in Table 3, with the Test Articles or a Reference compound (Ref. compound). The reference compound was provided by Jackson Laboratory and the identity remained confidential. Levamisole was provided in a 550 mg tablet (Allay Pharmaceuticals LLC) and prepared fresh at each use+6. Thymalfasin (T5) was provided as 1.6 mg lyophilized powder in a 5 ml vial for reconstitution with sterile water.

**Table 3.** Experimental cohorts and study design

| Group | # Male mice per study arm | Genotype | Treatment | Dosing Route | Dosing Frequency |
| --- | --- | --- | --- | --- | --- |
| 1 | 12 | nontransgenic | Vehicle | PO | QD for 2 weeks then QOD for 6 weeks |
| 2 | 18 | G93A-SOD1 | Vehicle |  |  |
| 3 | 18 | G93A-SOD1 | Reference Cpd 20 mg/kg |  |  |
| 4 | 18 | G93A-SOD1 | Test Article 1 (levamisole) 1 mg/kg |  |  |
| 5 | 18 | G93A-SOD1 | Test Article 2 (thymosin α1) 0.04 mg/kg | SQ | 2x/week for 8w |
| 1 | 12 | nontransgenic | Vehicle | PO | QD for 2 weeks then QOD for 6 weeks |
| 2 | 18 | TDP43-A315T | Vehicle |  |  |
| 3 | 18 | TDP43-A315T | Reference Cpd 20 mg/kg |  |  |
| 4 | 18 | TDP43-A315T | Test Article 1 (levamisole) 1 mg/kg |  |  |
| 5 | 18 | TDP43-A315T | Test Article 2 (thymosin α1) 0.04 mg/kg | SQ | 2x/week for 8w |

### Measured parameters

Body weights were measured 3 days prior to dosing, then twice a week at the time of dosing. Mice were assessed for overall body condition weekly from 8 to 14 weeks of age, and daily thereafter. At 12 weeks of age (after 4 weeks of treatment), latency to fall on the wire hang test was measured in all mice of Groups 1-5 in the TDP43-A315T model. At 10 and 12 weeks of age (after 2 and 4 weeks of treatment), Compound Muscle Action Potential (CMAP) was measured under anesthesia in mice of Groups 1-5 in the *B6SJL-Tg(SOD1*G93A)1Gur/J* model. The amplitude of the CMAP is reported for the maximal response recorded upon increasing stimulation intensities. The value is in mV and read peak-to-peak. Repetitive Nerve Stimulation (RNS) consists of 10 CMAPs recorded at a supramaximal stimulation intensity; stimulation is made of 10 successive pulses at 3, 6, 10 and 20 Hz. The decrement is calculated as the % decrease of the CMAP amplitude between the first response and the lowest one in the train of 10. At 14 weeks of age (after 6 weeks of treatment), 12 mice in each group were humanely euthanized by CO2 narcosis. For up to 6 mice per group, the time to humane endpoint was quantified. No tissue was collected in mice reading end-stage.

### Neuropathological evaluations

The spinal cord, brain, femoral nerve and 6 other organs from all mice collected at 14 weeks, and the colon of all mice of Groups 1 to 5 collected at 14 weeks of age, were fixed for histology. Spinal cord and brain were paraffin-embedded, sectioned (three slides of 3 sections each per tissue and per mouse), and stained with anti-ubiquitin, GFAP and Iba1 IHC (one slide per stain). Immuno-reactivity was quantified on slide scans as the percentage of a region of interest (ROI) positive for the stain, over the area of the ROI (one ROI on the cortex, and one ROI on the spinal cord, per mouse). Femoral nerves were cross-sectioned, stained with Toluidine Blue, and the number and caliber distribution of the motor and sensory axons were quantified on one section per mouse (specific methods not supplied by Jackson Laboratories). The myenteric plexus of the colon was micro-dissected, stained for acetylcholinesterase activity and ganglion cells were quantified in one ROI per mouse.

### Statistical Methods

The statistical tests used are described in the legends of each figure as appropriate.

## Results

For each model, we generated multiple cohorts to build towards a target number of 18 animals per treatment group with 12 nontransgenic controls. At 8 weeks of age, treatments were initiated and mice were treated for up to 8 weeks as described in Table 3. Mice were treated with levamisole by oral gavage, while Thymosin α1 was delivered by s.c. injection following the schedules outlined in Table 3. Mice were also treated with a proprietary reference compound for comparison. For each genotype and treatment group, a random subset of 6 animals were allowed to age to humane endpoints to assess drug effects on survival. The remaining 12 animals were euthanized after 8 weeks of treatment for neuropathological analyses.

### B6SJL-Tg(SOD1*G93A)1Gur/J Model

None of the treatments significantly improved survival or mitigated weight-loss in G93A SOD1 mice (Fig. 1A, B). At 10 and 12 weeks of age, we analyzed muscle function using electrophysiological stimulation. At 10 weeks of age (2 weeks of treatment), the compound action potential in treated and untreated G93A mice was lower than non-transgenic controls (Fig. 2A). At 12 weeks of (4 week of treatment), compound muscle action potential declined further with no evidence that any treatment mitigated loss of muscle function (Fig. 2B). We also conducted repetitive nerve stimulation to assess muscle function at 10 and 12 weeks of age. Compared to nontransgenic controls, G93A SOD1 mice showed reduced compound action potentials at both 10 and 12 weeks of age (Fig 3 A-H). None of the treatments consistently improved muscle function. Mice treated with Thymosin α1 trended towards diminished function relative to G93A mice treated with vehicle (Fig. 3A-H). As expected for the G93A model, there were reduced numbers of motor axons by 16 weeks of age (Fig. 4A). None of the treatments resulted in increased motor axons counts (Fig. 4A). Sensory axon numbers were not significantly diminished in 16 month-old G93A mice, and none of the treatments altered sensory axon numbers (Fig. 4B). At 16 weeks of age, the spinal cords of vehicle-treated G93A mice had higher levels of GFAP, Iba1, and ubiquitin immunoreactivity (Fig. 5A-C). None of the treatments lowered GFAP or Iba1 reactivity (Fig. 5A, B). Intriguingly, all three treatments reduced the burden of ubiquitin immunoreactivity (Fig. 5C). Collectively, these data indicate that neither levamisole nor thymosin α1 treatment produced significant improvements in survival, muscle, function, or motor neuron loss. The only observed positive effect was a reduction in ubiquitin immunoreactivity in the treated mice.

**Figure 1.**
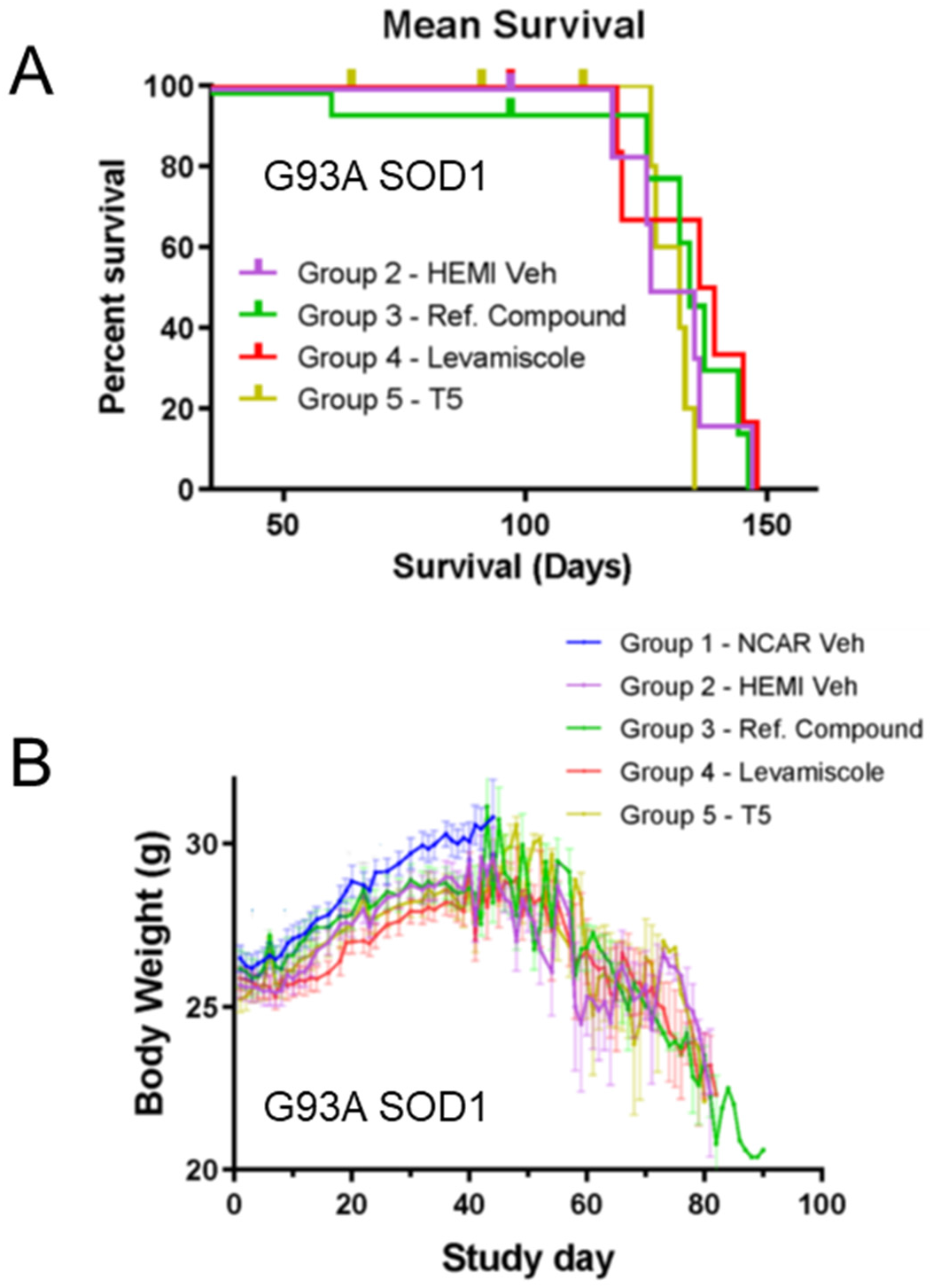
Survival and weight data for G93A SOD1 mice treated with reference compound, levamisole, and thymosin α1. G93A SOD1 mice were generated in two cohorts with a random subset of six mice from each treatment group allowed to age to end-stage paralysis. None of the treatments prolonged survival. Weights were measured as described in Methods beginning at 8 weeks of age and through the course of the dosing study (n = 18 per treatment group for G93A SOD1 mice; n = 12 for nontransgenic controls). All treatment groups reached peak weight at about the same age before declining. At 8 weeks after treatment, all nontransgenic controls and 12 G93A SOD1 mice from each treatment group were euthanized for tissue collection. No differences in weight gain or retention by any treatment were observed.. NCAR = nontransgenic control. Hemi = hemizygous transgene positive. T5 = thymosin α1.

**Figure 2.**
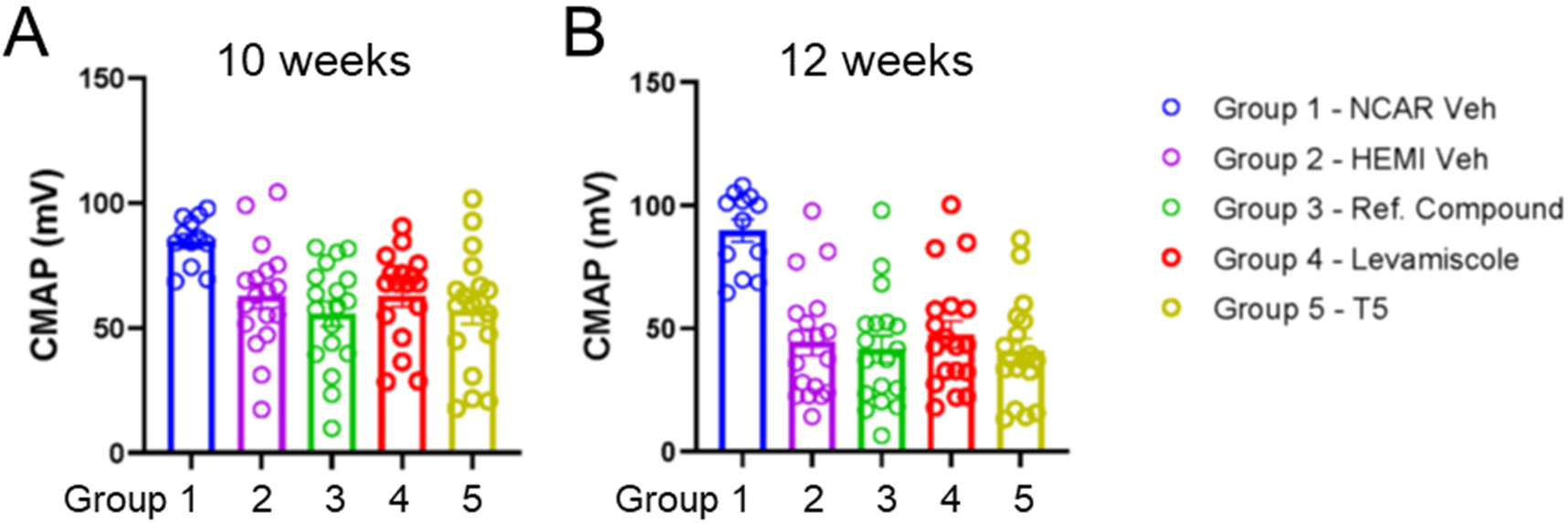
Compound action potential measurements in G93A SOD1 mice treated with reference compound, levamisole, and thymosin α1. A. By 10 weeks of age (2 weeks of treatment), measurements detected reduced amplitude compound action potentials in vehicle-treated G93A SOD1 mice relative to vehicle-treated nontransgenic controls (unpaired two-tailed T-test Group 1 vs Group 2; p 0.0023). None of the treatment groups were statistically different from vehicle treated G93A SOD1 mice (one way ANOVA Group 2 vs Groups 3-5; F 0.5368; p 0.6587). B. At 12 weeks of age (4 weeks of treatment), measurements again showed a reduction in the amplitude of compound action potentials in vehicle-treated G93A SOD1 mice relative to vehicle-treated nontransgenic mice (unpaired two-tailed T-test Group 1 vs Group 2; p 0.0001). None of the treatments mitigated the reduction in compound action potential amplitudes (one way ANOVA Group 2 vs Groups 3-5; F 0.3310; p 0.8029). n = 15-18 per treatment group for G93A SOD1 mice; n = 12 for nontransgenic controls)

**Figure 3.**
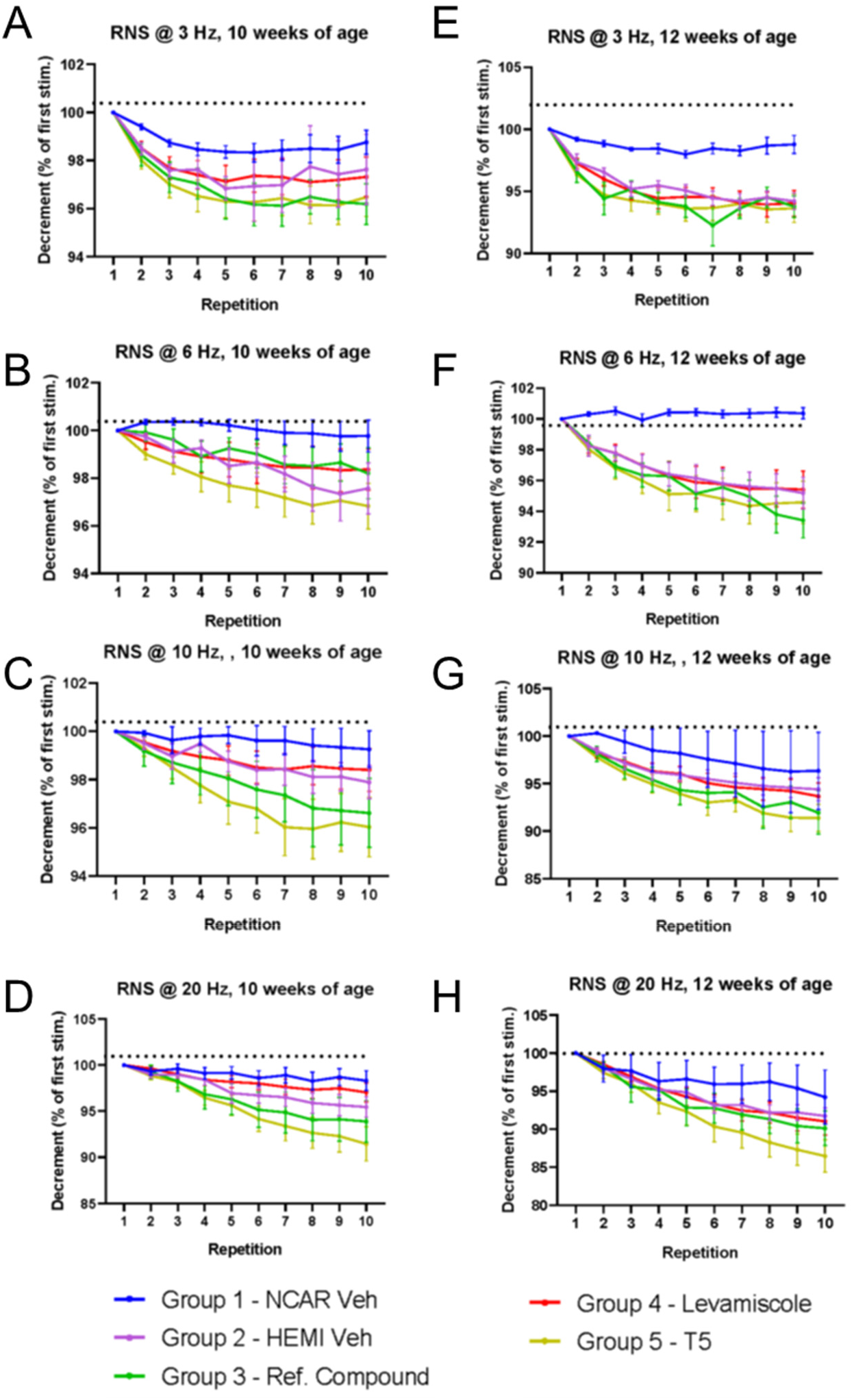
Measurements of the reduction in compound action potentials after repeated nerve stimulation at escalating frequencies. A-D. Reductions in compound action potential relative to the first stimulation for each treatment group are graphed for repeated nerve stimulation measurements at 10 weeks of age (2 weeks of treatment). E-H. Graphs for data obtained at 12 weeks of age (4 weeks of treatment). At both time points, G93A SOD1 mice treated with thymosin α1 tended to show the greatest reductions in compound action potentials after repeated stimulation.

**Figure 4.**
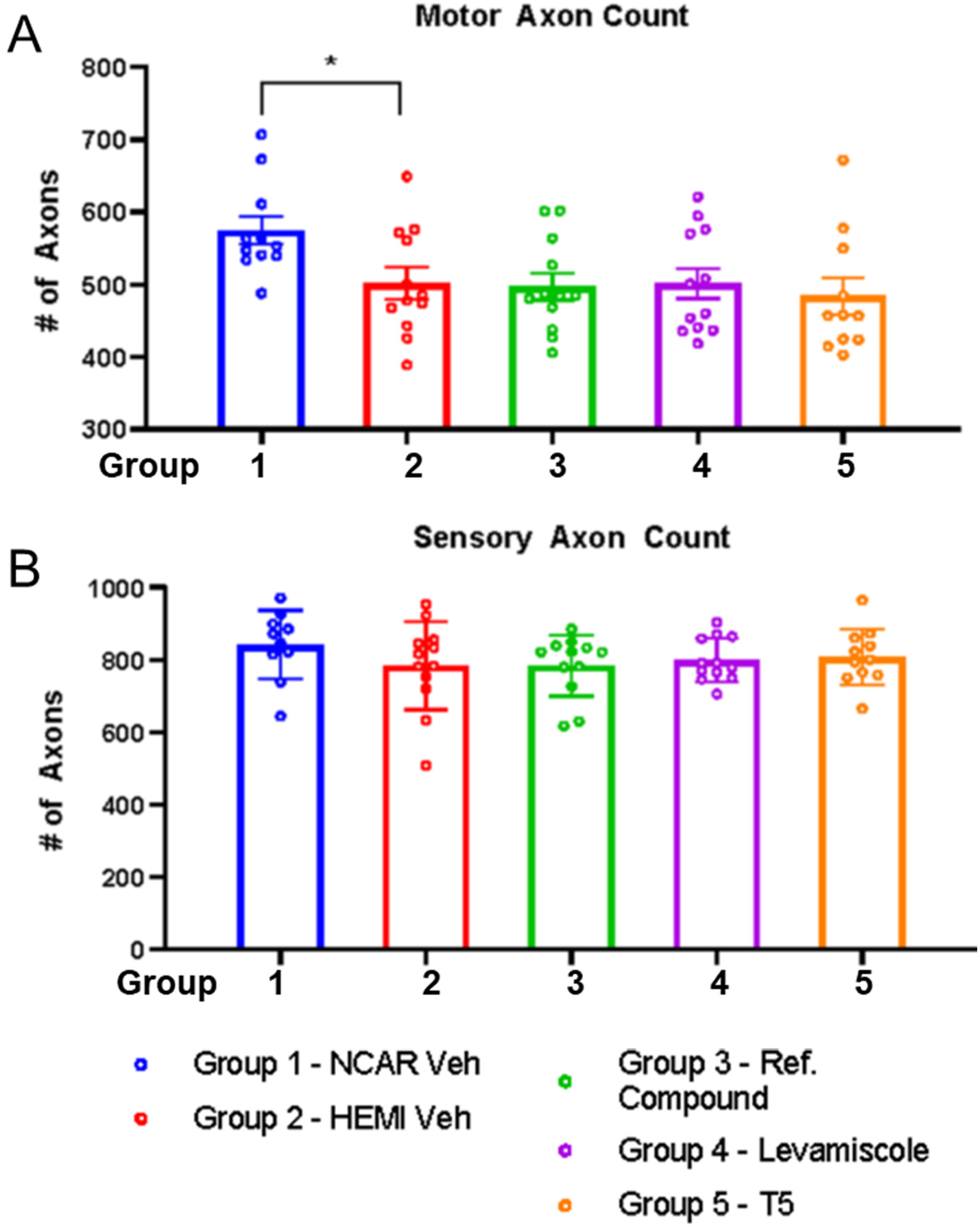
Selective loss of motor axons in femoral nerves that is not mitigated by any treatment tested. A. Data for motor axon counts from the femoral nerve of 16 week old mice in each treatment group (n = 11-12 per genotype per group). Vehicle-treated G93A SOD1 mice showed reduced numbers of motor axons relative to vehicle-treated nontransgenic controls (unpaired two-tailed T-test Group 1 vs Group 2; p 0.0205). None of the treatment groups were statistically different from vehicle treated G93A SOD1 mice (one way ANOVA Group 2 vs Groups 3-5; F 0.1474; p 0.9308). B. Data for sensory axons counts from the femoral nerves at 16 weeks of age. The number of sensory axons in vehicle-treated G93A SOD1 mice was similar to vehicle-treated nontransgenic controls (unpaired two-tailed T-test Group 1 vs Group 2; p 0.2362). None of the treatment groups were statistically different from vehicle treated G93A SOD1 mice (one way ANOVA Group 2 vs Groups 3-5; F 0.21; p 0.8889). n = 12 per group per genotype.

**Figure 5.**
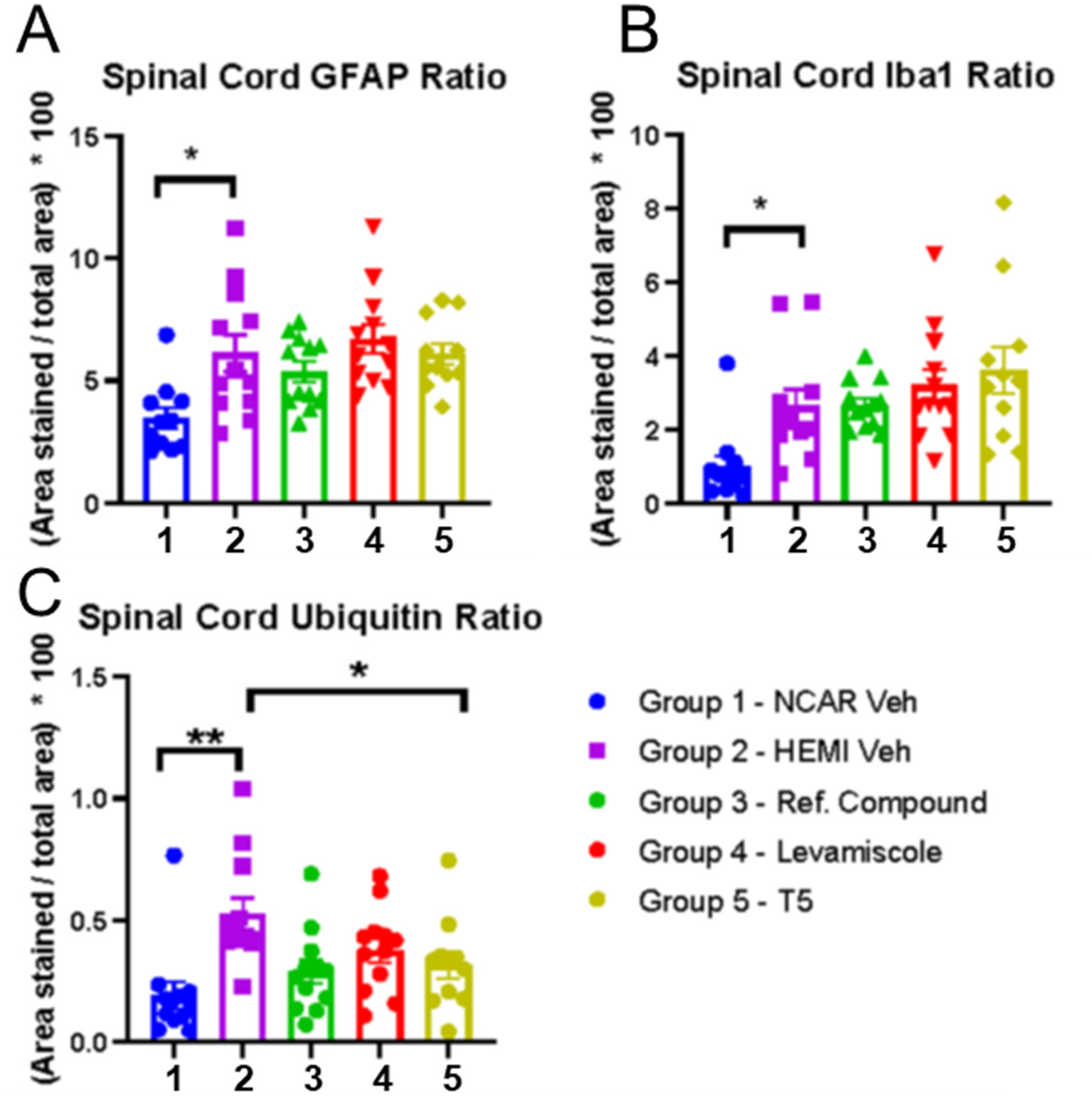
Analysis of GFAP, Iba-1, and ubiquitin immunoreactivity in the spinal cords of G93A SOD1 mice from all treatment groups. A. Analysis of GFAP immunoreactivity graphed as area stained/total area. The fractional area of GFAP immunoreactivity in vehicle-treated G93A SOD1 mice was higher than vehicle-treated nontransgenic controls (unpaired two-tailed T-test Group 1 vs Group 2; p 0.069). GFAP burden in spinal cords of G93A SOD1 mice from the treatment groups was statistically similar to vehicle treated G93A SOD1 mice (one way ANOVA Group 2 vs Groups 3-5; F 0.9682; p 0.4165). B. Analysis of Iba-1 immunoreactivity graphed as fractional area. The fractional area of Iba-1 immunoreactivity in vehicle-treated G93A SOD1 mice was higher than vehicle-treated nontransgenic controls (unpaired two-tailed T-test Group 1 vs Group 2; p 0.0073). Iba-1 burden in spinal cords of G93A SOD1 mice from the treatment groups was similar to vehicle treated G93A SOD1 mice (one way ANOVA Group 2 vs Groups 3-5; F 1.067; p 0.3733). C. Analysis of ubiquitin immunoreactivity graphed as fractional area. The fractional area of ubiquitin immunoreactivity in vehicle-treated G93A SOD1 mice was higher than vehicle-treated nontransgenic controls (unpaired two-tailed T-test Group 1 vs Group 2; p 0.0007). Ubiquitin burden in spinal cords of G93A SOD1 mice from the treatment groups was lower than that of vehicle treated G93A SOD1 mice (one way ANOVA Group 2 vs Groups 3-5; F 3.791; p 0.0169). n = 12 per group per genotype.

### B6.Cg-Tg(Prnp-TARDBP*A315T)95Balo/J Model

Treatment with levamisole or thymosin α1 did not significantly extend the survival or mitigate weight-loss in TDP43-A315T mice (Fig. 6A, B). To examine muscle function, TDP43-A315T mice were examined on a wire hanging test, measuring latency to fall at 12 weeks of age. Vehicle treated TDP43-A315T mice showed a latency to fall that was not significantly different from nontransgenic controls (Fig. 7). Mice in all three treatment groups showed similar levels of performance on the wire hanging task (Fig. 7). At 16 weeks of age, the spinal cords of vehicle treated TDP43-A315T mice had higher levels of GFAP and Iba1 immunoreactivity (Fig. 8A, B). The levels of these markers in all three treatment groups were similar to vehicle-treated mice (Fig. 8A, B). The levels of ubiquitin immunoreactivity in the spinal cord of vehicle-treated TDP43-A315T mice was similar to nontransgenic controls (Fig. 8C). Mice treated with the reference compound and levamisole trended towards higher levels of ubiquitin immunoreactivity, but the differences did not reach statistical significance.

**Figure 6.**
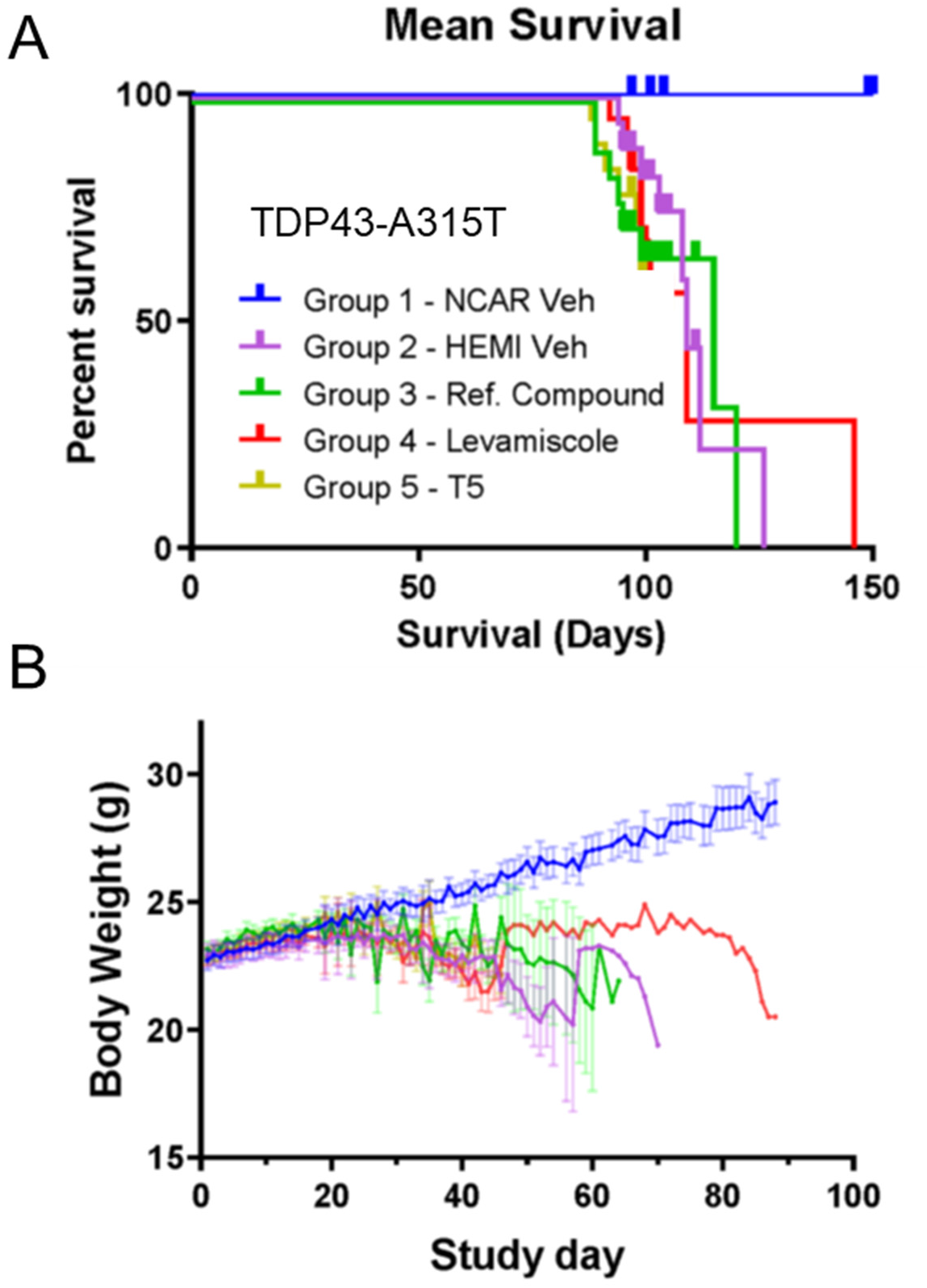
Survival and weight data for TDP43-A315T mice treated with reference compound, levamisole, and thymosin α1. A. TDP43-A315T mice were generated in three cohorts with a random subset of six mice from each treatment group allowed to age to humane end-points. None of the treatments dramatically prolonged survival. B. Weights were measured as described in Methods beginning at 8 weeks of age and through the course of the dosing study (n = 18 per treatment group for TDP43-A315T mice; n = 12 for nontransgenic controls). All treatment groups reached peak weight at about the same age before declining. At 8 weeks after treatment, all nontransgenic controls and 12 TDP43-A315T mice from each treatment group were euthanized for tissue collection. No differences in weight gain or retention by any treatment were observed. The color tracings in B are the same legend as in panel A.

**Figure 7.**
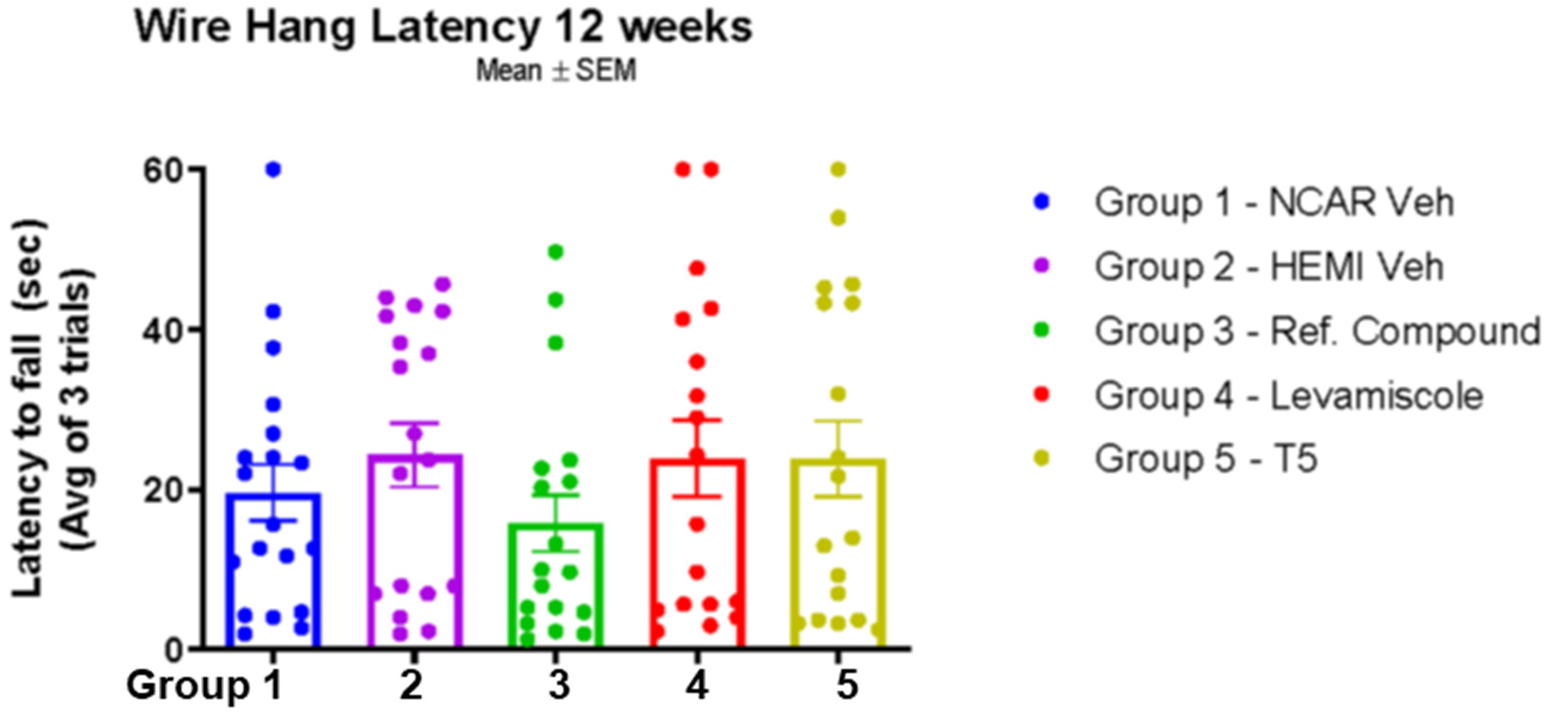
Assessment of grip strength by wire hanging task in TDP43-A315T mice. The latency for mice to fall from suspended wire was assessed as described in Methods. There was no difference in latency to fall between vehicle-treated nontransgenic mice and vehicle treated TDP-43-A315T mice (unpaired two-tailed T-test Group 1 vs Group 2; p 0.3768). There was no difference in performance between vehicle-treated TDP-43-A315T mice and any of the treatment groups (one way ANOVA Group 2 vs Groups 3-5; F 0.9230; p 0.4345). n=18-19 per genotype per treatment group.

**Figure 8.**
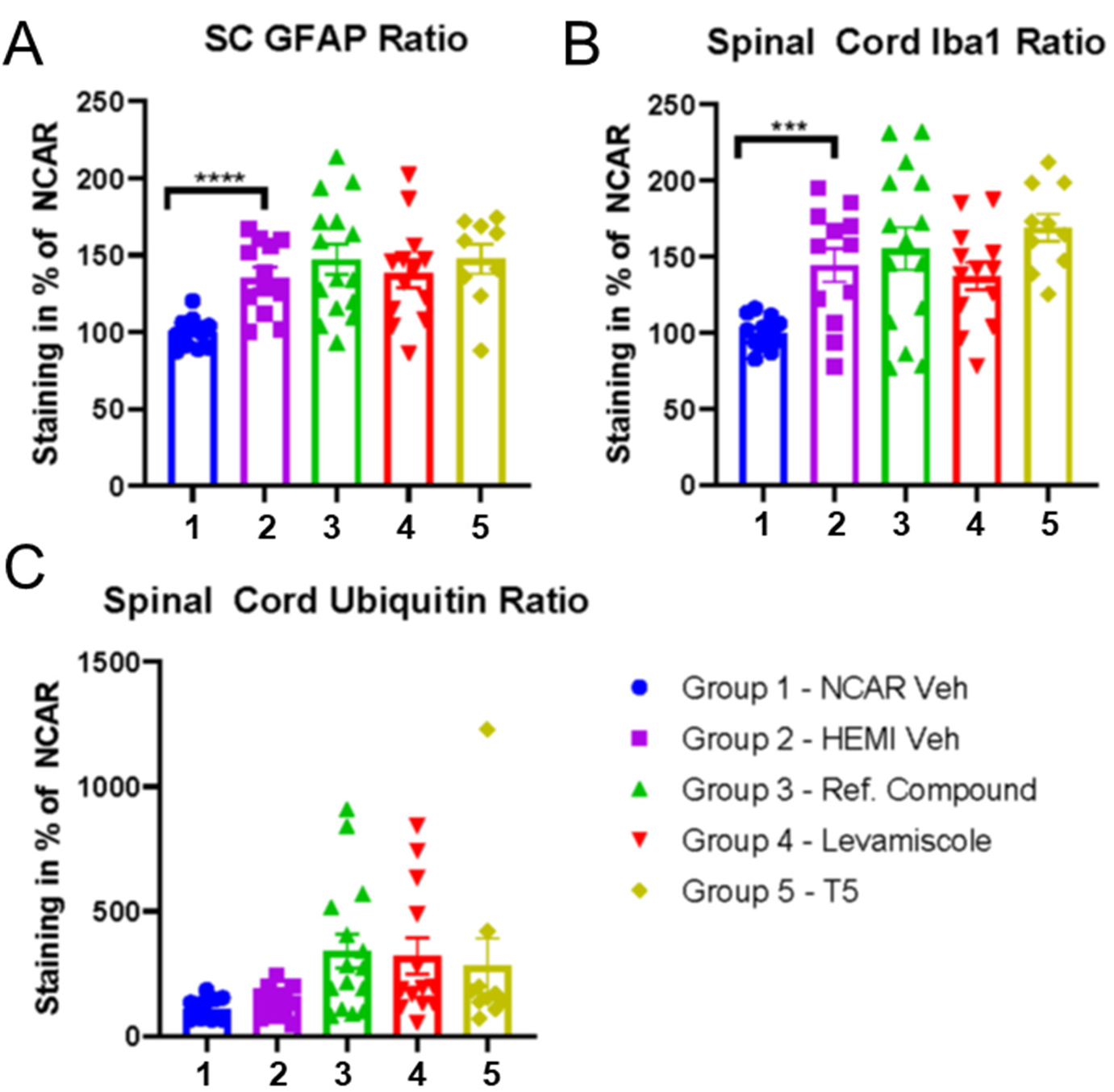
Analysis of GFAP, Iba-1, and ubiquitin immunoreactivity in the spinal cords of TDP43-A315T mice from all treatment groups. A. Analysis of GFAP immunoreactivity graphed as area stained/total area. The fractional area of GFAP immunoreactivity in vehicle-treated TDP43-A315T mice was higher than vehicle-treated nontransgenic controls (unpaired two-tailed T-test Group 1 vs Group 2; p 0.0001). GFAP burden in spinal cords of TDP43-A315T mice from the treatment groups was similar to vehicle treated TDP43-A315T mice (one way ANOVA Group 2 vs Groups 3-5; F 0.4783; p 0.6990). B. Analysis of Iba-1 immunoreactivity graphed as fractional area. The fractional area of Iba-1 immunoreactivity in vehicle-treated TDP43-A315T mice was higher than vehicle-treated nontransgenic controls (unpaired two-tailed T-test Group 1 vs Group 2; p 0.0008). Iba-1 burden in spinal cords of TDP43-A315T mice from the treatment groups was similar to vehicle treated TDP43-A315T mice (one way ANOVA Group 2 vs Groups 3-5; F 1.319; p 0.2796). C. Analysis of ubiquitin immunoreactivity graphed as fractional area. The fractional area of ubiquitin immunoreactivity in vehicle-treated TDP43-A315T mice was similar to vehicle-treated nontransgenic controls (unpaired two-tailed T-test Group 1 vs Group 2; p 0.1438). Ubiquitin burden in spinal cords of TDP43-A315T mice from the treatment groups was not different from vehicle treated TDP43-A315T mice (one way ANOVA Group 2 vs Groups 3-5; F 1.658; p 0.1891). n = 9-15 per group per genotype.

## Discussion

The current studies were designed to the effects of levamisole and Thymosin α1 on pathogenic processes in ALS, with a small cohort of 6 mice per treatment group used to assess effects on lifespan. The B6/SJL G93A SOD1 model is a reliable mimic of motor neuron disease [4] but the TDP43-A315T mice in congenic C57BL6/J backgrounds used here are confounded by gastrointestinal dysfunction that is largely driving abbreviated lifespan [7,8]. At the ages used in this study, the motor deficits in TDP43-A315T mice were mild and the only pathologic abnormalities were increased neuroinflammatory markers. None of the treatments significantly improved measures of muscle function in either model. Survival was increased slightly in the levamisole treated cohorts in the TDP43 model, but neither drug extended survival in the G93A SOD1 model. Although both of the drugs tested here are described as immune modulators, neither drug treatment led to significant reductions in inflammatory pathological markers in either model. The only statistically significant pathologic finding was that all drugs tested, including the reference compound, reduced the burden of ubiquitin immunostaining in the G93A SOD1 model.

Thymosin α1 is an immune modulator proposed for treating rheumatoid arthritis and that has some effects in the central nervous system [12–14]. The full-length peptide is 49 amino acids long, but pentapeptide corresponding to amino acids 32-36 accounts for all of its biological activities [15]. Thymalfasin (T5) is a pro-molecule of thymopoietin that has been shown to affect the function of T cells and monocytes [16,17]. Thymosin α1 has also been described as an anti-inflammatory molecule. It has been reported to suppress NF-kB activation and inhibit microglial activation by reducing secretion of inflammatory mediators [18]. Although these studies indicate that Thymosin α1 could modulate immune reactions in various disease settings, we found no evidence that our treatment paradigm influenced neuroinflammatory markers in either the G93A SOD1 or TDP43 model.

Thymosin α1 was initially described as binding the acetyl choline receptor (AchR) in a manner that interferes with αBgt binding sites and decreases neuromuscular transmission [19]. Subsequently, investigators determined that Thymopoeitin preparations used in these studies were contaminated with α-cobratoxin or phospholipase A_2_, and concluded that these contaminates may have been responsible for the activities attributed to thymosin α1. Work by Ochoa and colleagues examined the activities of synthetic thymosin α1 in AchR binding studies, finding little or no direct binding [20]. Instead, these investigators found evidence that thymosin α1 accelerates the desensitization of AchR, which could diminish neuromuscular transmission. Notably, in our repeated nerve stimulation studies of muscle function in G93A SOD1 mice, we observed that mice treated with thymosin α1 frequently showed the largest decrement in compound action potentials (see Fig. 3). These findings suggest that some activities of thymosin α1 could be a therapeutic liability in the treatment of ALS.

Levamisole is an active agonist for subset of nicotinic AchRs, reportedly stimulating muscle contraction with a primary action on parasitic nematode (Ca^++^) ion channels [21,22]. In mammals, levamisole can inhibit or facilitate ganglionic Ach receptors and these actions may enhance neuromuscular function [21]. It was anticipated that levamisole could affect compound action potential measurements because it affects noradrenergic transmission in the peripheral nervous system by inhibiting noradrenergic reuptake. However, there was no significant treatment effect on compound action potentials in G93A SOD1 mice, nor was there consistent effects of levamisole on repeated nerve-stimulation assessments of muscle function. Overall, we found no evidence that levamisole treatment at the doses used here produced any detectable benefit in muscle function.

The pharmacokinetics of levamisole in humans has been extensively studied [23–25]. Levamisole has been reported to inhibit tissue non-specific alkaline phosphatase (TNAP) in brain blood vessels to reversibly increase blood brain barrier permeability [26]. Levamisole HCl reduces neuronal response amplitudes and decreases axonal conduction by blocking voltage-dependent sodium channels [22]. The effects of blocking voltage-dependent sodium channels may be suppression of axonal growth, myelination, and synaptic plasticity [22]. Our study in these two mouse models did not reveal any deleterious effects of levamisole at the dose used here.

Levamisole has strong immunomodulatory activity [27–29]. Early in its development large-scale clinical research programs tested the effect of levamisole in human diseases caused by impaired cellular immune mechanisms, including ALS where the drug was ineffective [11]. More promising results were obtained in a small trial of multiple sclerosis patients [30]. Levamisole has dose dependent actions that may not be beneficial to patients. Individuals exposed to levamisole through illicit drugs exhibit many adverse reactions [10]. If levamisole were to be developed for treatment of neurological diseases a significant consideration would be preventing the reversible side effects through drug design [31].

Collectively, we observe that treatment with levamisole or thymosin α1 produced little benefit in G93A SOD1 or TDP43-A315T mice. Treatment effects of the Reference Compound were not superior to levamisole or thymosin α1 in either model. The only positive indication we detected for any of the tested drugs was a reduction in ubiquitin load in the spinal cord of G93A SOD1 mice. The effects of these drugs on ubiquitin were mild in this model with levamisole only trending towards significance. The potential mechanism by which these drugs could interfere with ubiquitin accumulation is unknown. Modifications to levamisole or thymosin α1 could heighten immune modulation in a manner that would be beneficial to ALS patients. This study demonstrates that these drugs have no adverse impact in two mouse models. Adding these drugs in combination with other medications could produce synergistic benefits that would prolong the survival of ALS patients.

## Acknowledgements

We thank Laurent Bogdanik (Jackson Laboratories, Bar Harbor, ME) for assistance in executing this project. This work was entirely funded by Neurodegenerative Disease Research, Inc., a non-profit entity supporting research in ALS.

